# Loading of piperlongumine to liposomes after complexation with ß- cyclodextrin and its effect on viability of colon and prostate cancer cells

**DOI:** 10.1101/314161

**Authors:** Kalana W. Jayawardana, Nidhi Jyotsana, Zhenjiang Zhang, Michael R. King

**Affiliations:** Vanderbilt University, Department of Biomedical Engineering, 5824 Stevenson Center, Nashville, TN 37235 USA

**Author notes:** These authors contributed equally (Kalana W. Jayawardana, Nidhi Jyotsana). The authors declare no conflict of interest.

**Keywords:** Piperlongumine, Liposomes, ß-Cyclodextrin, COLO-205 cells, PC-3 cells

## Abstract

Use of nano carriers to treat cancer is attractive due to their advantages such as the sustained release of drugs and ability to target specific regions of the body where treatment is needed. However, loading water insoluble chemotherapeutic drugs into liposomes is challenging. In this study, we developed a method to encapsulate water-insoluble drug (piperlongumine) in liposomes by complexing piperlongumine with ß-Cyclodextrin. Liposomes encapsulated with piperlongumine incubated with COLO 205 and PC-3 cell lines and demonstrated that viability of COLO 205 and PC-3 cells decreases to 7% and 41% respectively when the piperlongumine concentration is at 20 μM.

## Introduction

Currently, there is great interest in loading drugs into nanocarriers to create controlled release drug carriers with pulsatile release kinetics.^1^ In recent times cancer therapeutics have been explored for delivery via nanocarriers. Use of this technology for cancer treatment has several advantages. Some of these advantages include the fact that drugs delivered using nanocarriers can maintain the therapeutic concentration for extended periods compared to free drug, and the fact that nanocarriers can be used to target specific areas of the body where tumors are present and increase the drug concentration in the area of interest.^2^ For these reasons, much of the current work is devoted to the development of new materials to produce nanocarriers to load water insoluble drugs that are difficult to load into conventional nanocarriers such as liposomes. However, the use of liposomes as nanocarriers has its own advantages. For example, liposomes have been extensively tested for human use and approved by the United States Food and Drug Administration (FDA). Liposomes are considered safe for human consumption because of their biocompatible lipid matrix. Another advantage is that liposomes can be engineered to have different sizes and different permeability properties, therefore, release kinetics can be controlled depending on specific requirements.^3–4^

Drug loading into liposomes is achieved by either passive loading or active loading.^5–6^ In passive loading, dried lipid film is hydrated by an aqueous solution containing the drug. In active loading, drug is internalized to preformed liposomes by introducing a transmembrane pH gradient.^7^ The passive loading approach can only be used for drugs that are water soluble. Most of the drugs used for chemotherapy have poor water solubility, therefore, it is challenging to load drugs into liposomes using passive delivery. This motivates the development of drug formulations to load water-insoluble drugs into liposomes.

Cyclodextrins (CD) are a family of cyclic sugars that have a hydrophilic outer surface and hydrophobic inner cavity. CD molecules are relatively large with a number of hydrogen donors and acceptors. Because of the presence of hydrophobic cavity in CD, hydrophobic drugs can be incorporated into this cavity and bind to form stable complexes, but allow slow efflux of the CD-entrapped drugs to be released once administered in vivo.^8^ Therefore, CD formulation of water-insoluble drugs increases their water solubility and bioavailability. Furthermore, CDs are nontoxic, biologically inert, and currently, there are over 30 products worldwide containing drug/CD based formulations on the market.^9^ Previous work from our lab has demonstrated the TRAIL enhanced piperlongumine (PL) induced apoptosis of colon and prostate cancer cell lines.^10^ Therefore, in this study we developed a CD and piperlongumine formulation in PBS buffer to increase the water solubility of the piperlongumine, and used this solution to engineer liposomes to interact with the colon cancer cell line COLO 205 and prostate cancer cell line PC-3.

## Results and Discussion

PL aqueous solution was prepared by dropwise addition of tetra hydro furan solution of PL into a preheated aqueous solution of ß-CD. Liposomes used in this study were prepared by drying a lipid formulation consisting of L-α phosphatidylcholine (Egg PC), cholesterol, and DSPE-PEG under a vacuum desiccator followed by hydration of the dried lipid film with an aqueous solution of CD and PL (Figure 1). This mixture was then extruded through a membrane with pore size similar to the desired particle size. In this study, we used a membrane with pore size diameter 100 nm. Liposome sizes were measured to be 112 nm ± 53 nm (DLS) (Figure 2). The resulting liposomes were dialyzed to remove excess piperlongumine before performing *in vitro* studies.

**Figure 1.**
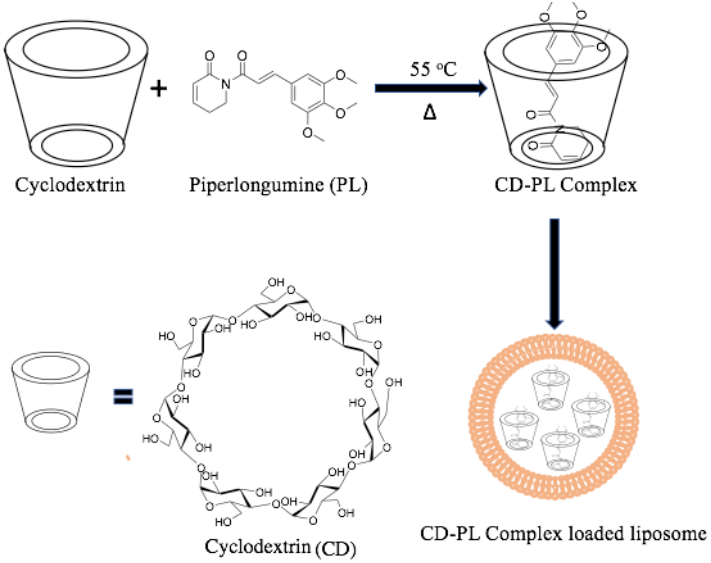
Schematic representation of CD-PL complex formation and liposome formulation

**Figure 2.**
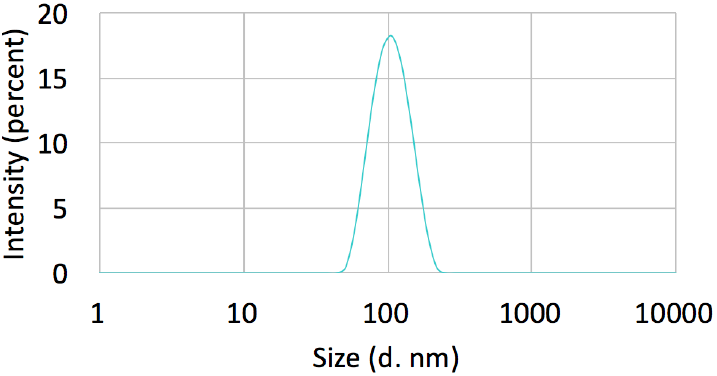
DLS graph of piperlongumine liposomes

*In vitro* studies were performed by incubating PL liposomes with the colon cancer cell line COLO 205 and the prostate cancer cell line PC-3. Control studies were performed by incubating cells with naked CD liposomes. COLO 205 and PC-3 cancer cells were treated with different concentrations of CD-PL or naked liposomes (N.CD). Following 24 hr of treatment, AlamarBlue cell viability and Annexin V apoptosis assays were performed. Both COLO 205 and PC-3 cells remained unaffected by naked liposomes, however cell viability decreased with increasing PL concentrations (Figure 3). COLO 205 cells responded significantly better to PL treatment compared to PC-3 cells. At 20 μM and 40 μM PL concentrations, the viability of COLO 205 cells decreased to 7% and 5%. PC-3 cells decreased to 41% and 7%, respectively.

**Figure 3.**
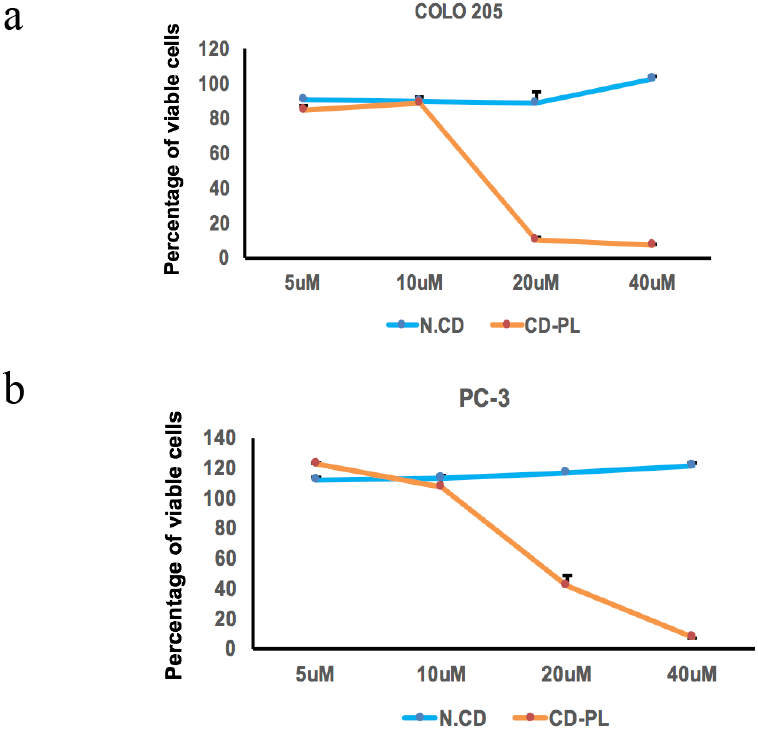
Viability of (a) COLO 205 and (b) PC-3 cells were analyzed by AlamarBlue assay at 24 hr after treatment with cyclodextrin-piperlongumine at different concentrations (5, 10, 20 and 40 μM piperlongumine) (mean ± SEM, n=3). Viability data for naked liposomes shown in blue, CD-PL liposomes shown in green.

Similar trends were observed using the flow cytometry-based Annexin-V apoptosis assay. Almost all COLO 205 cells underwent apoptosis at higher PL concentrations, whereas 50% of PC-3 cells were apoptotic at the highest PL concentration (Figure 4).

**Figure 4.**
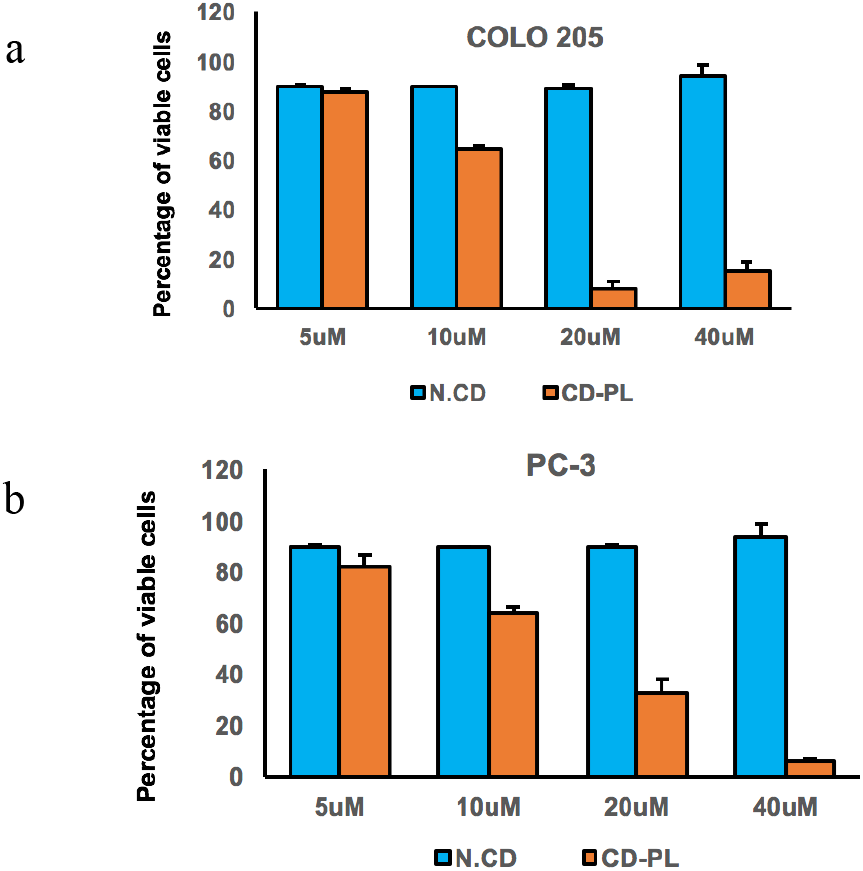
Percent of viable (a) COLO 205 and (b) PC-3 cells analyzed by Annexin V assay at 24 hr after treatment with cyclodextrin-piperlongumine (CD-PL) at different concentrations (5, 10, 20 and 40 μM PL) (mean ± SEM, n=2).

In summary, we have demonstrated that CD can enhance the water solubility of PL. PL is a water-insoluble drug, but with complexation with CD, we can achieve increase solubility of PL in aqueous media. Furthermore, we have demonstrated that we can engineer liposomes consisting of CD-PL complex and it can be used to kill COLO 205 cells *in vitro*.

## Materials and Methods

### Increase of piperlongumine concentration in PBS buffer with CD

Cyclodextrin (20 mg/mL, 5 mL) in PBS buffer was heated to 55 OC and piperlongumine (5 mg/mL, 0.5 mL) in tetrahydrofuran (THF) was added dropwise. The mixture was stirred for 24 hr and cooled to room temperature. The solution was filtered through a 100 μm filter. Before making liposomes, the solution was filtered through the liposome membrane (100 nm), and lipid film was hydrated using this filtered CD-PL solution.

### Loading of CD-PL into liposomes

Volumes of 616 μL of Egg PC, 155 μL of cholesterol, and 337 μL of DSPE-PEG (Avanti Polar Lipids) were mixed in a small vial and the mixture dried under vacuum to form a thin lipid film. Then the lipid film was hydrated with a CD-PL-containing PBS buffer solution and sonicated for 30 min. The resulting drug-encapsulated liposome solution was extruded with a 100 nm filter 30 times. The resulting liposome solution was dialyzed with fresh PBS solution for 24 hr.

### Cell Lines and Cell Culture

The colon cancer cell line COLO 205 (ATCC number CCL-222) was obtained from ATCC and cultured in RPMI 1640 supplemented with 2 mM L-Glutamine, 25 mM Hepes, 10% (vol/vol) FBS and 100 U/mL PenStrep (complete media) under humidified conditions at 37°C and 5% CO_2_. The prostate cancer cell line PC-3 (ATCC number CRL-1435) was obtained from ATCC and cultured in F-12K medium supplemented with 10% (vol/vol) FBS and 100 U/mL PenStrep.

### Incubation of CD-piperlongumine liposomes with COLO 205 and PC-3 cells

COLO 205 and PC-3 cells were seeded in multi-well plates at a seeding density of 100,000 cells/mL, 1 d before experimentation to ensure that the cells were in the linear phase of the growth cycle. The cell media was changed before experimentation. Cells were incubated with either the solvent 1X DPBS or cyclodextrin-piperlongumine. The treated cells were maintained in culture conditions for 24 hr and later analyzed by Annexin-V assay and AlamarBlue assay to quantify the proportion of viable cells.

### Flow cytometry experiment to determine cell viability and degree of apoptosis

Quantitation of cell death was analyzed using the Annexin-V apoptosis assay on a Guava easyCyte™ flow cytometer (Millipore, Billerica, MA, USA). Samples were prepared per the manufacturer’s instructions. Cells were classified into four categories based on dye uptake: viable cells (negative for Annexin-V and propidium iodide (PI)), early apoptotic cells (positive for Annexin-V only), late apoptotic cells (positive for Annexin-V and PI), and necrotic cells (positive for PI only).

### AlamarBlue Assay

10,000 cells were seeded in 100 μl of media in each well of a 96-well flat bottom transparent plate. 1/10th volume of the AlamarBlue^®^ reagent (786-921, G-Biosciences) was directly added to the wells and incubated for 4 h at 37°C in a cell culture incubator, protected from direct light. Results were recorded by measuring fluorescence using a fluorescence excitation wavelength of peak excitation 570 nm and peak emission 585 nm on a microplate reader (Tecan Infinite 500, Tecan Group Ltd).

## End Matter

### Author Contributions and Notes

The manuscript was written through contributions of all authors. All authors have given approval to the final version of the manuscript.

## Acknowledgments

This work was supported by National Institute of Health Grant No. R01 CA203991 to M.R.K.

## References

1. Hubbell, J. A.; Langer, R., Translating materials design to the clinic. Nature Materials 2013, 12, 963.

2. Kohay, H.; Sarisozen, C.; Sawant, R.; Jhaveri, A.; Torchilin, V. P.; Mishael, Y. G., PEG-PE/clay composite carriers for doxorubicin: Effect of composite structure on release, cell interaction and cytotoxicity. Acta Biomaterialia 2017, 55, 443–454.

3. Blok, M. C.; Van Der Neut-Kok, E. C. M.; Van Deenen, L. L. M.; De Gier, J., The effect of chain length and lipid phase transitions on the selective permeability properties of liposomes. Biochimica et Biophysica Acta (BBA) - Biomembranes 1975, 406 (2), 187–196.

4. Koyanagi, T.; Leriche, G.; Onofrei, D.; Holland, G. P.; Mayer, M.; Yang, J., Cyclohexane Rings Reduce Membrane Permeability to Small Ions in Archaea - Inspired Tetraether Lipids. Angewandte Chemie International Edition 2016, 55 (5), 1890–1893.

5. Kita, K.; Dittrich, C., Drug delivery vehicles with improved encapsulation efficiency: taking advantage of specific drug–carrier interactions. Expert Opinion on Drug Delivery 2011, 8 (3), 329–342.

6. Schwendener, R. A.; Schott, H., Liposome Formulations of Hydrophobic Drugs. In Liposomes: Methods and Protocols, Volume 1: Pharmaceutical Nanocarriers, Weissig, V., Ed. Humana Press: Totowa, NJ, 2010; pp 129–138.

7. Abraham, S. A.; Edwards, K.; Karlsson, G.; Hudon, N.; Mayer, L. D.; Bally, M. B., An evaluation of transmembrane ion gradient-mediated encapsulation of topotecan within liposomes. Journal of Controlled Release 2004, 96 (3), 449–461.

8. Szejtli, J., Introduction and General Overview of Cyclodextrin Chemistry. Chemical Reviews 1998, 98 (5), 1743–1754.

9. Loftsson, T.; Jarho, P.; Másson, M.; Järvinen, T., Cyclodextrins in drug delivery. Expert Opinion on Drug Delivery 2005, 2 (2), 335–351.

10. Li, J.; Sharkey, C. C.; King, M. R., Piperlongumine and immunecytokine TRAIL synergize to promote tumor death. Scientific Reports 2015, 5, 9987

